## Supplemental File 2 for "A Timeline of Bacterial and Archaeal Diversification in the Ocean"

((((RS.GCF.006274605.1.protein:2739.07000,
(GB.GCA.001587575.1.protein:2205.88000,GB.GCA.004212155.1.protein:2205
.88000)Node_1065:533.18400)Node_1064:355.58700,
((RS.GCF.000018485.1.protein:1726.79000,RS.GCF.000025525.1.protein:172
6.79000)Node_1067:1190.81000,(RS.GCF.000166095.1.protein:1900.51000,
(GB.GCA.003584625.1.protein:1130.17000,
(GB.GCA.001316325.1.protein:794.88500,RS.GCF.000828575.1.protein:794.8
8500)Node_1070:335.28800)Node_1069:770.33800)Node_1068:1017.09000)Node
_1066:177.05700)Node_1063:146.85500,
((((GB.GCA.002067865.1.protein:1451.98000,RS.GCF.000404225.1.protein:1
451.98000)Node_1060:900.20900,(RS.GCF.000025665.1.protein:1839.38000,
(RS.GCF.900176435.1.protein:788.58700,RS.GCF.000011185.1.protein:788.5
8700)Node_1062:1050.79000)Node_1061:512.80400)Node_1059:208.59500,
((GB.GCA.002498725.1.protein:726.47100,
(GB.GCA.002501805.1.protein:571.90900,
(GB.GCA.002503285.1.protein:462.37200,GB.GCA.002504905.1.protein:462.3
7200)Node_1054:109.53700)Node_1053:154.56100)Node_1052:511.21600,
(GB.GCA.002720095.2.protein:1037.73000,
(GB.GCA.003602415.1.protein:670.20000,
(GB.GCA.002722135.1.protein:539.51700,
(GB.GCA.002499195.1.protein:410.60700,GB.GCA.002494645.1.protein:410.6
0700)Node_1058:128.91100)Node_1057:130.68300)Node_1056:367.53200)Node_
1055:199.95400)Node_1051:1323.09000)Node_1050:408.60400,
((GB.GCA.002011165.1.protein:2407.25000,
(RS.GCF.000025505.1.protein:1845.97000,GB.GCA.002010305.1.protein:1845
.97000)Node_1049:561.28000)Node_1048:238.34800,
(GB.GCA.004212075.1.protein:2497.97000,
((GB.GCA.001766815.1.protein:2132.55000,
(RS.GCF.000063445.1.protein:1981.61000,
(RS.GCF.000235565.1.protein:1785.78000,
(RS.GCF.000685155.1.protein:1444.41000,GB.GCA.004212095.1.protein:1444
.41000)Node_1047:341.37200)Node_1046:195.82500)Node_1045:150.94400)Nod
e_1044:154.61800,
((RS.GCF.000021965.1.protein:945.91400,RS.GCF.001571405.1.protein:945.
91400)Node_10401328514.35:1111.77000,
(GB.GCA.002503845.1.protein:1017.66000,
(RS.GCF.000334895.1.protein:580.07000,
(RS.GCF.000230735.2.protein:410.52900,RS.GCF.900215575.1.protein:410.5
2900)Node_1043:169.54100)Node_1042:437.58900)Node_1041:1040.03000)Node
_1039:229.48200)Node_1038:210.79900)Node_1037:147.63000)Node_1036:323.
78700)Node_1035:272.12600)Node_1034:250.70000,
(((GB.GCA.003144275.1.protein:2955.31000,GB.GCA.002728275.1.protein:29
55.31000)Node_1033:328.35200,((GB.GCA.001940655.1.protein:2188.14000,
(GB.GCA.004524545.1.protein:2017.83000,GB.GCA.004524535.1.protein:2017
.83000)Node_1032:170.30900)Node_1031:921.14800,
(GB.GCA.004524565.1.protein:3027.12000,
(GB.GCA.001940665.1.protein:2809.63000,
(GB.GCA.005191415.1.protein:2199.31000,GB.GCA.005191425.1.protein:2199
.31000)Node_1030:610.31900)Node_1029:217.48800)Node_1028:82.17490)Node
_10273238.54:174.37400)Node_1026:172.40400,

(((RS.GCF.000011205.1.protein:2586.86000,
(RS.GCF.000092185.1.protein:2336.20000,
(RS.GCF.000017945.1.protein:2021.90000,RS.GCF.000223395.1.protein:2021
.90000)Node_102222381793.31:314.30100)Node_1021:250.66000)Node_1020:47
1.11100,
((GB.GCA.003649445.1.protein:2345.50000,GB.GCA.003649495.1.protein:234
5.50000)Node_1024:391.16100,
(GB.GCA.002506515.1.protein:1737.33000,GB.GCA.000015805.1.protein:1737
.33000)Node_1025:999.33500)Node_1023:321.30800)Node_1019:236.55000,
((GB.GCA.005888735.1.protein:2778.41000,RS.GCF.003589585.1.protein:277
8.41000)Node_1018:306.07700,
(((GB.GCA.000270325.1.protein:1866.06000,GB.GCA.003056285.1.protein:18
66.06000)Node_1017:324.34600,
(GB.GCA.002898395.1.protein:1647.19000,GB.GCA.000494145.1.protein:1647
.19000)Node_1016:543.22300)Node_1015:799.96800,
(GB.GCA.002499005.1.protein:2578.20000,
((GB.GCA.005877225.1.protein:2030.56000,
(GB.GCA.002495315.1.protein:1823.94000,GB.GCA.002495905.1.protein:1823
.94000)Node_1014:206.61700)Node_1013:298.47200,
(GB.GCA.900248165.1.protein:1961.26000,
(GB.GCA.002713325.1.protein:1676.22000,
((GB.GCA.000802205.2.protein:1197.43000,
(RS.GCF.000698785.1.protein:1015.03000,
(GB.GCA.005877075.1.protein:871.24900,GB.GCA.003176995.1.protein:871.2
4900)Node_1012:143.78200)Node_1011:182.39500)Node_1010:350.76600,
((GB.GCA.900177045.1.protein:745.26300,
(GB.GCA.007280335.1.protein:537.17900,GB.GCA.013330055.1.protein:537.1
7900)Node_1009:208.08500)Node_1008:257.95800,
((RS.GCF.002787055.1.protein:227.97500,GB.GCA.002499525.1.protein:227.
97500)Node_996:585.02000,(GB.GCA.001443365.1.protein:674.57300,
(((GCA.902592435.1.AG-447-
C19.genomic:143.00600,GB.GCA.001627235.1.protein:143.00600)Node_1005:3
59.67200,(73669.assembled:80.75760,
(13609.assembled:62.03910,2513237066:62.03910)Node_1007:18.71850)Node_
1006:421.92000)Node_1004:100.12800,
(GB.GCA.000200715.1.protein:526.73200,
((RS.GCF.000220175.1.protein:120.62800,GB.GCA.000204585.1.protein:120.
62800)Node_1003:208.97000,(GB.GCA.000299365.1.protein:198.49400,
(RS.GCF.000956175.1.protein:80.74880,RS.GCF.003175215.1.protein:80.748
80)Node_1002:117.74500)Node_1001:131.10500)Node_1000:197.13300)Node_99
9:76.07290)Node_998:71.76730)Node_997:138.42300)Node_995:190.22600)Nod
e_994:544.97000)Node_993:128.02800)Node_992:285.04100)Node_991:367.769
00)Node_990:249.16900)Node_989:412.18000)Node_988:94.10850)Node_987:21
0.03600)Node_986:161.54600)Node_985:36.14070)Node_984:1164.31000,
((GB.GCA.003641555.1.protein:3592.59000,
(RS.GCF.000745455.1.protein:3133.99000,
((RS.GCF.000504085.1.protein:2105.00000,RS.GCF.001990485.1.protein:210
5.00000)Node_983:421.77200,(GB.GCA.003538775.1.protein:2355.21000,
(RS.GCF.000953715.1.protein:1949.11000,RS.GCF.002752635.1.protein:1949
.11000)Node_982:406.09400)Node_981:171.56600)Node_980:607.21800)Node_9

79:458.60200)Node_978:626.07900,
((GB.GCA.001443005.1.protein:3847.94000,
((GB.GCA.001508935.1.protein:2125.80000,RS.GCF.900129645.1.protein:212
5.80000)Node_977:143.17300,(RS.GCF.000233775.1.protein:1987.29000,
((RS.GCF.000025885.1.protein:1472.03000,RS.GCF.000237805.1.protein:147
2.03000)Node_976:322.27800,(RS.GCF.000526375.1.protein:1572.05000,
(GB.GCA.002432945.1.protein:1377.02000,RS.GCF.001701045.1.protein:1377
.02000)Node_975:195.03800)Node_974:222.25400)Node_973:192.98000)Node_9
72:281.68700)Node_971:1578.97000)Node_970:360.58500,
((((RS.GCF.000184705.1.protein:3673.79000,
((RS.GCF.000621445.1.protein:1376.09000,
(RS.GCF.003815035.1.protein:1041.75000,
(RS.GCF.000798955.1.protein:896.20900,
(RS.GCF.000313915.1.protein:354.06200,RS.GCF.000392875.1.protein:354.0
6200)Node_962:542.14700)Node_961:145.54100)Node_960:334.34200)Node_959
:1727.99000,
((GB.GCA.002407935.1.protein:2676.15000,GB.GCA.002411145.1.protein:267
6.15000)Node_969:319.35200,
((RS.GCF.000213255.1.protein:2122.30000,GB.GCA.002426575.1.protein:212
2.30000)Node_965:206.53200,(RS.GCF.900142285.1.protein:2112.79000,
(RS.GCF.002874775.1.protein:1479.97000,
(RS.GCF.900167005.1.protein:1292.80000,RS.GCF.003460745.1.protein:1292
.80000)Node_968:187.16700)Node_967:632.82200)Node_966:216.03800)Node_9
64:666.67100)Node_963:108.58400)Node_958:569.70300)Node_957:316.68100,
((((GB.GCA.003134215.1.protein:2727.81000,GB.GCA.002431715.1.protein:2
727.81000)Node_956:352.86100,(GB.GCA.002898555.1.protein:2600.76000,
(GB.GCA.003577165.1.protein:2006.13000,GB.GCA.002162095.1.protein:2006
.13000)Node_955:594.63400)Node_954:479.91200)Node_953:329.56000,
((GB.GCA.002898575.1.protein:3057.90000,
(RS.GCF.002973605.1.protein:2634.10000,GB.GCA.001872605.1.protein:2634
.10000)Node_952:423.80300)Node_951:213.44200,
(GB.GCA.002305165.1.protein:2912.87000,
(GB.GCA.003170595.1.protein:2071.85000,GB.GCA.003542955.1.protein:2071
.85000)Node_950:841.01700)Node_949:358.47600)Node_948:138.88800)Node_9
47:501.86800,
((RS.GCF.900175965.1.protein:3359.59000,GB.GCA.002305585.1.protein:335
9.59000)Node_932:88.05060,((RS.GCF.003386795.1.protein:1770.31000,
(RS.GCF.001660485.1.protein:1526.93000,
(RS.GCF.000701365.1.protein:1225.54000,RS.GCF.002926165.1.protein:1225
.54000)Node_946:301.39400)Node_945:243.38000)Node_944:1184.10000,
(((GB.GCA.002418265.1.protein:1157.34000,GB.GCA.002450985.1.protein:11
57.34000)Node_943:602.50200,(GB.GCA.002897675.1.protein:1557.77000,
(GB.GCA.003388545.1.protein:957.12000,GB.GCA.003242515.1.protein:957.1
2000)Node_942:600.64700)Node_941:202.07100)Node_940:851.03600,
(GB.GCA.002352645.1.protein:2368.01000,
((RS.GCF.000949295.1.protein:1283.02000,GB.GCA.003151475.1.protein:128
3.02000)Node_939:334.51800,(GB.GCA.004366205.1.protein:1367.48000,
(GCA.902570085.1.AG-414-
N20.genomic:544.67500,GB.GCA.002687745.1.protein:544.67500)Node_938:82
2.80400)Node_937:250.06200)Node_936:750.46500)Node_935:242.86900)Node_

934:343.53700)Node_933:493.22700)Node_931:464.46300)Node_930:78.36840)
Node_929:54.96420,
((((GB.GCA.002747975.1.protein:1196.07000,RS.GCF.002761215.1.protein:1
196.07000)Node_928:1791.29000,(GB.GCA.001818315.1.protein:2615.70000,
(GB.GCA.003551435.1.protein:2032.36000,
(GB.GCA.001189155.1.protein:963.12100,GB.GCA.003504275.1.protein:963.1
2100)Node_927:1069.24000)Node_926:583.34100)Node_925:371.65800)Node_92
4:845.86500,((GB.GCA.001898225.1.protein:3379.56000,
(GB.GCA.004295045.1.protein:2791.17000,
(GB.GCA.001567485.1.protein:2075.92000,GB.GCA.001796335.1.protein:2075
.92000)Node_923:715.24600)Node_922:588.38900)Node_921:87.81110,
(GB.GCA.004377275.1.protein:3055.65000,
(GB.GCA.003523055.1.protein:2457.83000,((GCA.902588795.1.AG-439-
D18.genomic:2238.29000,GB.GCA.003228195.2.protein:2238.29000)Node_920:
139.49400,
((GB.GCA.002711005.1.protein:1644.97000,GB.GCA.002719675.1.protein:164
4.97000)Node_919:400.94200,((GB.GCA.002238855.1.protein:1646.20000,
(GB.GCA.002313895.1.protein:1014.50000,GB.GCA.002328865.1.protein:1014
.50000)Node_918:631.70800)Node_917:211.77400,
(GB.GCA.002721445.1.protein:1570.30000,
(GB.GCA.002696685.1.protein:890.29100,
(GB.GCA.002717465.1.protein:749.18000,(GCA.902578465.1.AG-426-
M11.genomic:660.02400,GB.GCA.002728195.1.protein:660.02400)Node_916:89
.15670)Node_915:141.11100)Node_914:680.01100)Node_913:287.67400)Node_9
12:187.93500)Node_911:331.87700)Node_910:80.04140)Node_909:597.82100)N
ode_908:411.71700)Node_907:365.85400)Node_906:170.90200,
((((GB.GCA.001771535.1.protein:2102.18000,GB.GCA.001771585.1.protein:2
102.18000)Node_905:471.14900,
(GB.GCA.001771545.1.protein:2139.63000,GB.GCA.001771575.1.protein:2139
.63000)Node_904:433.70000)Node_903:866.17100,
((GB.GCA.003242895.1.protein:2889.54000,GB.GCA.003864475.1.protein:288
9.54000)Node_902:126.98900,
((GB.GCA.003265975.1.protein:1804.63000,GB.GCA.003265885.1.protein:180
4.63000)Node_900:385.14700,
(GB.GCA.002433595.1.protein:1993.43000,GB.GCA.002296285.1.protein:1993
.43000)Node_901:196.34700)Node_899:826.74500)Node_898:422.97700)Node_8
97:339.46700,((GB.GCA.003503675.1.protein:3341.88000,
(GB.GCA.005110385.1.protein:2554.72000,GB.GCA.001899315.1.protein:2554
.72000)Node_859:787.15700)Node_858:136.61800,
(RS.GCF.000011385.1.protein:2690.67000,
(RS.GCF.000013205.1.protein:2237.63000,
((GB.GCA.003242085.1.protein:408.20400,GB.GCA.003249035.1.protein:408.
20400)Node_896:1516.77000,(RS.GCF.002754935.1.protein:1799.37000,
((RS.GCF.001939115.1.protein:1546.46000,
((RS.GCF.001870905.1.protein:1197.11000,RS.GCF.000014265.1.protein:119
7.11000)Node_895:137.94000,(RS.GCF.000317635.1.protein:1027.98000,
(GB.GCA.000252485.1.protein:847.77000,
(RS.GCF.000332035.1.protein:744.91000,
(RS.GCF.000167195.1.protein:215.76200,
(RS.GCF.000169335.1.protein:86.81450,RS.GCF.000017845.1.protein:86.814

50)Node_894:128.94700)Node_893:529.14800)Node_892:102.85900)Node_891:1
80.21200)Node_890:307.07000)Node_889:211.40400)Node_888:137.14000,
(RS.GCF.000010065.1.protein:1298.37000,
(GB.GCA.001007665.1.protein:671.94600,
(RS.GCF.003011125.1.protein:528.51700,
((RS.GCF.003011885.1.protein:331.39200,
(GB.GCA.000708525.1.protein:126.05400,
(RS.GCF.000316515.1.protein:17.41410,RS.GCF.003003925.1.protein:17.414
10)Node_887:108.64000)Node_886:205.33800)Node_885:98.67740,
((RS.GCF.000011485.1.protein:214.92600,
(RS.GCF.000018585.1.protein:136.70000,
((RS.GCF.000012465.1.protein:19.13060,GB.GCA.003279655.1.protein:19.13
060)Node_876:84.11830,(RS.GCF.000015665.1.protein:20.96900,
(RS.GCF.000012645.1.protein:8.44350,
(RS.GCF.000015965.1.protein:4.77420,RS.GCF.000018065.1.protein:4.77420
)Node_875:3.66930)Node_874:12.52550)Node_873:82.27990)Node_872:33.4511
0)Node_871:78.22620)Node_870:48.67780,
(((RS.GCF.004209775.1.protein:128.37800,RS.GCF.000153825.1.protein:128
.37800)Node_880:41.26860,
(RS.GCF.000014585.1.protein:141.28400,RS.GCF.000063505.1.protein:141.2
8400)Node_879:28.36230)Node_878:54.89610,
(RS.GCF.000195975.1.protein:125.24600,
(RS.GCF.000153805.1.protein:96.75660,
(RS.GCF.000737595.1.protein:22.80130,
(RS.GCF.000012625.1.protein:13.24870,RS.GCF.900474085.1.protein:13.248
70)Node_884:9.55260)Node_883:73.95520)Node_882:28.48930)Node_881:99.29
660)Node_877:39.06160)Node_869:166.46500)Node_868:98.44740)Node_867:14
3.42900)Node_866:626.42600)Node_865:385.22400)Node_864:115.77700)Node_
863:125.60000)Node_862:312.65800)Node_861:453.04200)Node_860:787.82300
)Node_857:300.47400)Node_856:225.15500)Node_855:41.30970)Node_854:76.1
3320,(((GB.GCA.004297825.1.protein:3180.94000,
(GB.GCA.003695005.1.protein:2749.94000,
(RS.GCF.000017685.1.protein:2345.45000,RS.GCF.000243335.1.protein:2345
.45000)Node_853:404.49100)Node_852:430.99500)Node_851:403.32800,
((GB.GCA.001829155.1.protein:2988.28000,GB.GCA.002450905.1.protein:298
8.28000)Node_850:338.84000,(GB.GCA.001829125.1.protein:2866.93000,
(GB.GCA.002238925.1.protein:2248.85000,
((GB.GCA.002307015.1.protein:1495.10000,
(RS.GCF.000219725.1.protein:1305.85000,GB.GCA.001829185.1.protein:1305
.85000)Node_845:189.24900)Node_844:517.99200,
(RS.GCF.000758165.1.protein:1845.51000,
((GB.GCA.003641695.1.protein:1330.62000,RS.GCF.000378205.1.protein:133
0.62000)Node_848:167.51700,
(GB.GCA.007116075.1.protein:1205.22000,RS.GCF.000190435.1.protein:1205
.22000)Node_849:292.91400)Node_847:347.38000)Node_846:167.58300)Node_8
43:235.75700)Node_842:618.08100)Node_841:460.18900)Node_840:257.14300)
Node_839:387.88700,((((GB.GCA.003695705.1.protein:2802.56000,
((GB.GCA.002684655.1.protein:1966.88000,
(GB.GCA.002748355.1.protein:1281.94000,GB.GCA.002685965.1.protein:1281
.94000)Node_767:684.94000)Node_766:421.97600,

((GB.GCA.004295085.1.protein:1076.67000,
((GB.GCA.002705055.1.protein:758.59600,GB.GCA.002746235.1.protein:758.
59600)Node_760:173.71100,
(GB.GCA.003696625.1.protein:722.89900,GB.GCA.002500505.1.protein:722.8
9900)Node_759:209.40800)Node_758:144.35900)Node_757:1062.92000,
((GB.GCA.002683825.1.protein:1282.83000,GB.GCA.004296785.1.protein:128
2.83000)Node_762:156.79500,(GB.GCA.005799545.1.protein:1301.94000,
(GB.GCA.002686265.1.protein:1167.45000,
(GB.GCA.002686945.1.protein:979.70600,GB.GCA.002686595.1.protein:979.7
0600)Node_765:187.74500)Node_764:134.48600)Node_7631548.82:137.69100)N
ode_761:699.95900)Node_756:249.27400)Node_755:413.69900)Node_754:960.1
5200,
(((GB.GCA.002787155.1.protein:2265.34000,GB.GCA.001804285.1.protein:22
65.34000)Node_781:1142.47000,((GB.GCA.002787095.1.protein:3091.98000,
(GB.GCA.002771475.1.protein:2856.56000,
(GB.GCA.003141835.1.protein:2646.27000,
(GB.GCA.002780005.1.protein:2305.14000,GB.GCA.001804165.1.protein:2305
.14000)Node_780:341.12700)Node_779:210.29600)Node_778:235.41700)Node_7
77:70.08060,(GB.GCA.003645805.1.protein:2977.37000,
((GB.GCA.003644445.1.protein:1448.28000,GB.GCA.003154195.1.protein:144
8.28000)Node_773:1013.21000,
((GB.GCA.002796125.1.protein:2004.24000,GB.GCA.003598135.1.protein:200
4.24000)Node_775:284.23000,
(GB.GCA.002753745.1.protein:1242.34000,GB.GCA.003522725.1.protein:1242
.34000)Node_776:1046.13000)Node_774:173.02700)Node_772:515.87700)Node_
771:184.69200)Node_770:245.75500)Node_769:290.26800,
(RS.GCF.000204255.1.protein:3385.12000,
((GB.GCA.002405835.1.protein:2969.40000,
((RS.GCF.001017655.1.protein:2323.59000,GB.GCA.003566475.1.protein:232
3.59000)Node_794:275.86500,
(GB.GCA.002448475.1.protein:2200.22000,GB.GCA.001803315.1.protein:2200
.22000)Node_795:399.23300)Node_793:369.94400)Node_792:93.59880,
((GB.GCA.001872735.1.protein:1771.54000,GB.GCA.002500665.1.protein:177
1.54000)Node_791:1031.20000,
((GB.GCA.002422425.1.protein:2062.01000,GB.GCA.002452515.1.protein:206
2.01000)Node_790:605.79900,(GB.GCA.003695015.1.protein:2049.73000,
((GB.GCA.002690655.1.protein:1516.05000,GB.GCA.002715965.1.protein:151
6.05000)Node_788:252.81000,
(GB.GCA.003219345.1.protein:1629.93000,GB.GCA.005809105.1.protein:1629
.93000)Node_789:138.92800)Node_787:280.87600)Node_786:618.07600)Node_7
85:134.93100)Node_784:260.25500)Node_783:322.12100)Node_782:312.96800)
Node_768:64.62680)Node_753:119.43200,
((GB.GCA.002780205.1.protein:3015.46000,
(((GB.GCA.001871125.1.protein:1776.48000,GB.GCA.001800285.1.protein:17
76.48000)Node_802:882.46800,(GB.GCA.002841695.1.protein:2416.39000,
(RS.GCF.002355835.1.protein:1893.42000,GB.GCA.001800315.1.protein:1893
.42000)Node_801:522.97700)Node_800:242.56000)Node_799:172.39600,
((GB.GCA.003452935.1.protein:1797.02000,GB.GCA.001799575.1.protein:179
7.02000)Node_807:826.88900,
((GB.GCA.001800075.1.protein:2227.36000,GB.GCA.003140895.1.protein:222

7.36000)Node_805:227.36300,
(GB.GCA.001800095.1.protein:1532.54000,GB.GCA.001800135.1.protein:1532
.54000)Node_806:922.17900)Node_804:169.18900)Node_803:207.43500)Node_7
98:184.11200)Node_797:808.79000,
(((GB.GCA.001509335.1.protein:2560.29000,
(GB.GCA.003645825.1.protein:1678.55000,GB.GCA.002347855.1.protein:1678
.55000)Node_827:881.74100)Node_826:186.36800,
((GB.GCA.002689995.1.protein:1413.36000,
(GB.GCA.002717625.1.protein:1194.27000,
(GB.GCA.002705575.1.protein:1041.60000,GB.GCA.002703645.1.protein:1041
.60000)Node_838:152.66300)Node_837:219.09800)Node_836:826.67300,
((GB.GCA.002329125.1.protein:1151.07000,GB.GCA.002471705.1.protein:115
1.07000)Node_835:718.32100,((GCA.902538565.1.AG-899-
G06.genomic:185.16100,GCA.902629855.1.AH-324-
M03.genomic:185.16100)Node_831:1314.97000,
(GB.GCA.001577055.1.protein:1361.48000,
(GB.GCA.002701785.1.protein:1202.67000,(GCA.902615915.1.AG-892-
A09.genomic:708.15200,GB.GCA.002724735.1.protein:708.15200)Node_834:49
4.52100)Node_833:158.80400)Node_832:138.65300)Node_830:369.26100)Node_
829:370.64700)Node_828:506.62300)Node_825:638.56700,
((RS.GCF.001535965.1.protein:2910.52000,
(GB.GCA.001803165.1.protein:2662.89000,
(GB.GCA.001899175.1.protein:1565.95000,GB.GCA.004292595.1.protein:1565
.95000)Node_812:1096.94000)Node_811:247.62500)Node_810:188.58100,
((RS.GCF.003353065.1.protein:2241.70000,RS.GCF.003285105.1.protein:224
1.70000)Node_824:333.91700,((RS.GCF.000430505.1.protein:1340.04000,
(RS.GCF.000014145.1.protein:1123.41000,RS.GCF.001611675.1.protein:1123
.41000)Node_823:216.62800)Node_822:405.75000,
((RS.GCF.000275825.1.protein:1172.15000,GB.GCA.001897535.1.protein:117
2.15000)Node_821:363.19400,(GB.GCA.002483345.1.protein:1390.19000,
(GB.GCA.001769385.1.protein:1097.93000,
(GB.GCA.007119395.1.protein:977.63700,
(GB.GCA.002393195.1.protein:770.82300,
(GB.GCA.003851245.1.protein:521.19500,GB.GCA.900319235.1.protein:521.1
9500)Node_820:249.62800)Node_819:206.81400)Node_818:120.29100)Node_817
:292.26000)Node_816:145.15600)Node_815:210.44300)Node_814:829.83300)No
de_813:523.47700)Node_809:286.12900)Node_808:439.02300)Node_796:57.894
10)Node_752:53.10310,(((RS.GCF.003633715.1.protein:2889.78000,
(RS.GCF.000021545.1.protein:1712.06000,
(GB.GCA.003972855.1.protein:1189.51000,
(RS.GCF.003664005.1.protein:919.01900,RS.GCF.000020785.1.protein:919.0
1900)Node_751:270.49400)Node_750:522.54500)Node_749:1177.72000)Node_74
8:772.75500,(RS.GCF.000517565.1.protein:3174.11000,
(RS.GCF.000021725.1.protein:1486.44000,
(RS.GCF.000010325.1.protein:1116.72000,
(GB.GCA.003945285.1.protein:925.96300,
(((RS.GCF.900187115.1.protein:646.50200,GB.GCA.002424475.1.protein:646
.50200)Node_747:82.91880,
(RS.GCF.000147355.1.protein:560.60700,GB.GCA.000276965.1.protein:560.6
0700)Node_746:168.81400)Node_745:66.30880,

((RS.GCF.900637395.1.protein:532.83500,RS.GCF.000024885.1.protein:532.
83500)Node_744:172.83100,(RS.GCF.000259255.1.protein:611.42900,
(GB.GCA.003561815.1.protein:493.24400,GB.GCA.002428055.1.protein:493.2
4400)Node_743:118.18500)Node_742:94.23720)Node_741:90.06360)Node_740:1
30.23300)Node_739:190.75600)Node_738:369.72000)Node_73718101135.26:168
7.67000)Node_736:488.42900)Node_735:194.31700,
((GB.GCA.002402325.1.protein:3328.84000,
((RS.GCF.000428885.1.protein:2955.27000,GB.GCA.003650445.1.protein:295
5.27000)Node_734:299.07200,((GB.GCA.002898535.1.protein:2826.42000,
(GB.GCA.003141555.1.protein:1882.24000,GB.GCA.002010665.1.protein:1882
.24000)Node_733:944.18300)Node_732:291.19600,
((GB.GCA.005223145.1.protein:2787.51000,GB.GCA.002722645.1.protein:278
7.51000)Node_725:195.69000,(GB.GCA.003223145.1.protein:2799.39000,
((GB.GCA.001767495.1.protein:2310.29000,
(GB.GCA.003223555.1.protein:2084.27000,GB.GCA.004356105.1.protein:2084
.27000)Node_729:226.01900)Node_728:254.84600,
(GB.GCA.003225315.1.protein:1975.42000,
(GB.GCA.003140235.1.protein:1712.46000,GB.GCA.004299485.1.protein:1712
.46000)Node_731:262.96000)Node_730:589.72100)Node_727:234.24700)Node_7
26:183.81000)Node_724:134.42400)Node_723:136.72400)Node_722:74.50010)N
ode_721:378.07600,
((((GB.GCA.002238965.1.protein:434.67800,GB.GCA.002377645.1.protein:43
4.67800)Node_720:1069.79000,(GB.GCA.001803565.1.protein:1351.72000,
(GB.GCA.002500605.1.protein:1192.25000,
((GB.GCA.002787235.1.protein:498.51200,RS.GCF.000341545.2.protein:498.
51200)Node_719:615.33300,
(GB.GCA.001804915.1.protein:960.61300,GB.GCA.003229375.1.protein:960.6
1300)Node_718:153.23200)Node_717:78.40840)Node_716:159.46300)Node_848:
152.74800)Node_714:821.77900,
(((GB.GCA.002897915.1.protein:893.06900,GB.GCA.002011735.1.protein:893
.06900)Node_713:353.57200,(GB.GCA.003235715.1.protein:1185.43000,
((GB.GCA.002897855.1.protein:897.90400,GB.GCA.002011745.1.protein:897.
90400)Node_712:134.28200,(GB.GCA.001871685.1.protein:935.74600,
(RS.GCF.000020985.1.protein:731.84500,
(GB.GCA.001803635.1.protein:507.99200,GB.GCA.002753335.1.protein:507.9
9200)Node_711:223.85300)Node_710:203.90200)Node_709:96.43970)Node_708:
153.24000)Node_707:61.21410)Node_706:510.93800,
(RS.GCF.000299235.1.protein:1538.51000,
(GB.GCA.001805205.1.protein:1487.82000,
((GB.GCA.005239925.1.protein:937.23800,GB.GCA.004297235.1.protein:937.
23800)Node_705:360.16300,(GB.GCA.004298335.1.protein:924.81800,
(GB.GCA.003695915.1.protein:761.82400,
((GB.GCA.005877775.1.protein:468.37600,GB.GCA.004296885.1.protein:468.
37600)Node_704:86.21380,(RS.GCF.000196815.1.protein:367.53800,
(RS.GCF.001458695.1.protein:215.73800,
(RS.GCF.001458735.1.protein:104.59000,RS.GCF.001458775.1.protein:104.5
9000)Node_703:111.14800)Node_702:151.80000)Node_701:187.05200)Node_700
:207.23400)Node_699:162.99300)Node_698:372.58300)Node_697:190.41800)No
de_696:50.68830)Node_695:219.07100)Node_694:568.66500)Node_693:1086.96
000,

(((((RS.GCF.900142125.1.protein:1540.63000,RS.GCF.000012885.1.protein:
1540.63000)Node_658:1199.89000,(GB.GCA.002782605.1.protein:2559.78000,
(GB.GCA.001595385.3.protein:1829.41000,GB.GCA.001797495.1.protein:1829
.41000)Node_660:730.37300)Node_659:180.73700)Node_657:120.25400,
((RS.GCF.000092205.1.protein:2303.95000,
(RS.GCF.000174435.1.protein:2155.51000,
(RS.GCF.000217795.1.protein:1841.35000,GB.GCA.002419025.1.protein:1841
.35000)Node_664:314.15900)Node_663:148.43600)Node_662:184.68300,
(RS.GCF.000195295.1.protein:2400.14000,
(GB.GCA.001874005.1.protein:2123.90000,
(GB.GCA.002753225.1.protein:1309.85000,
(GB.GCA.000961655.1.protein:1109.00000,
(RS.GCF.000472805.1.protein:960.24500,GB.GCA.003973265.1.protein:960.2
4500)Node_669:148.75200)Node_668:200.84800)Node_667:814.05600)Node_666
:276.23500)Node_665:88.49480)Node_661:372.14000)Node_656:93.74520,
((((GB.GCA.002787535.1.protein:2352.67000,GB.GCA.002343185.1.protein:2
352.67000)Node_680:160.67300,
(RS.GCF.001748245.1.protein:1771.06000,GB.GCA.001907975.2.protein:1771
.06000)Node_1046:742.28700)Node_678:121.75600,
(GB.GCA.001797335.1.protein:2332.61000,
((GB.GCA.002778785.1.protein:1237.22000,
(RS.GCF.002208115.1.protein:999.17300,GB.GCA.002321895.1.protein:999.1
7300)Node_675:238.05000)Node_674:988.29100,
(GB.GCA.002786715.1.protein:1950.50000,
(GB.GCA.001769805.1.protein:1331.89000,GB.GCA.002387735.1.protein:1331
.89000)Node_677:618.60300)Node_676:275.01700)Node_673:107.09800)Node_6
72:302.49100)Node_671:172.56600,
((GB.GCA.002433485.1.protein:1933.06000,GB.GCA.002343455.1.protein:193
3.06000)Node_692:681.89100,((GB.GCA.002721815.1.protein:2294.96000,
(GB.GCA.007120035.1.protein:2128.96000,
(GB.GCA.002863065.1.protein:1767.46000,GB.GCA.006226565.1.protein:1767
.46000)Node_691:361.50300)Node_690:165.99400)Node_689:191.04700,
((GB.GCA.003242475.1.protein:1836.45000,
(RS.GCF.000022145.1.protein:1433.38000,GB.GCA.002408385.1.protein:1433
.38000)Node_685:403.07100)Node_684:498.10100,
((GB.GCA.003158455.1.protein:1911.14000,GB.GCA.001464385.1.protein:191
1.14000)Node_688:253.37300,
(RS.GCF.001931505.1.protein:1723.96000,GB.GCA.002320815.1.protein:1723
.96000)Node_687:440.55600)Node_686:170.03800)Node_683:151.45400)Node_6
82:128.94900)Node_681:192.71600)Node_1064:146.84700)Node_6553099.62:13
8.79400,((GB.GCA.002683655.1.protein:1851.28000,
(GB.GCA.002726945.1.protein:1470.87000,
((GB.GCA.001469005.2.protein:333.81600,GB.GCA.002683785.1.protein:333.
81600)Node_654:953.93800,(GB.GCA.001627865.1.protein:606.49200,
(GB.GCA.002082305.1.protein:442.90600,GB.GCA.002685535.1.protein:442.9
0600)Node_653:163.58600)Node_652:681.26300)Node_651:183.11800)Node_650
:380.40700)Node_649:1127.46000,
((((GB.GCA.003233235.1.protein:1042.76000,GB.GCA.002779605.1.protein:1
042.76000)Node_585:693.01300,
((GB.GCA.002791545.1.protein:1184.46000,RS.GCF.000744415.1.protein:118

4.46000)Node_587:226.91900,
(RS.GCF.005818865.1.protein:952.67300,RS.GCF.003665155.1.protein:952.6
7300)Node_588:458.70900)Node_586:324.38900)Node_584:949.48500,
(RS.GCF.000175575.2.protein:2188.45000,
(RS.GCF.000011705.1.protein:1968.43000,
((RS.GCF.003333315.1.protein:1592.49000,GB.GCA.002420425.1.protein:159
2.49000)Node_583:275.26900,(GB.GCA.003251535.1.protein:1672.69000,
((RS.GCF.000828835.1.protein:1364.95000,
(GB.GCA.003250975.1.protein:1145.61000,GB.GCA.002170535.1.protein:1145
.61000)Node_582:219.34000)Node_581:199.47400,
(((RS.GCF.004364935.1.protein:1090.41000,RS.GCF.000772705.2.protein:10
90.41000)Node_580:252.06600,(GB.GCA.000531125.1.protein:1265.23000,
(RS.GCF.004684205.1.protein:1141.83000,
(GB.GCA.001914695.1.protein:859.23400,GB.GCA.003567475.1.protein:859.2
3400)Node_579:282.59600)Node_578:123.40500)Node_577:77.24360)Node_576:
161.94900,(GB.GCA.003696905.1.protein:1368.72000,
(((GB.GCA.002101225.1.protein:1065.35000,GB.GCA.002733715.1.protein:10
65.35000)Node_575:86.95500,
(RS.GCF.900180345.1.protein:877.27600,RS.GCF.004001325.1.protein:877.2
7600)Node_574:275.02900)Node_573:113.94500,
((RS.GCF.002211785.1.protein:957.92500,RS.GCF.000297215.2.protein:957.
92500)Node_572:180.67000,(RS.GCF.001558495.2.protein:1043.42000,
(RS.GCF.001457615.1.protein:887.29000,(((GCA.902593675.1.AG-447-
I15.genomic:187.73400,(GCA.902594515.1.AG-447-
K20.genomic:133.88100,GCA.902600705.1.AG-917-
C02.genomic:133.88100)Node_571:53.85350)Node_570:309.32100,
(GCA.902582615.1.AG-912-C22.genomic:361.50900,(GCA.902616715.1.AG-892-
C21.genomic:235.42900,GCA.902594855.1.AG-915-
K19.genomic:235.42900)Node_569:126.08000)Node_568:135.54700)Node_567:2
71.73900,(GB.GCA.002469845.1.protein:412.26100,
(GB.GCA.002378645.1.protein:280.49700,
((RS.GCF.000511875.1.protein:138.84900,GB.GCA.001438605.1.protein:138.
84900)Node_566:73.43060,(GB.GCA.002335945.1.protein:173.57700,
(GB.GCA.004211905.1.protein:107.15300,GB.GCA.002382445.1.protein:107.1
5300)Node_565:66.42420)Node_564:38.70210)Node_563:68.21710)Node_562:13
1.76400)Node_561:356.53300)Node_560:118.49600)Node_559:156.13300)Node_
558:95.17260)Node_5571258.11:127.65500)Node_556:102.46800)Node_555:135
.71000)Node_554:59.99330)Node_553:108.26900)Node_552:195.07100)Node_55
1:100.67200)Node_550:220.01700)Node_549:496.80700)Node_548:78.42080,
((RS.GCF.002109495.1.protein:2109.71000,
(GB.GCA.002753665.1.protein:1826.27000,
(GB.GCA.002753565.1.protein:1469.64000,
(RS.GCF.004217665.1.protein:280.85400,GB.GCA.002753135.1.protein:280.8
5400)Node_648:1188.78000)Node_647:356.63700)Node_646:283.43300)Node_64
5:548.55300,(GB.GCA.003450915.1.protein:2381.93000,
((GB.GCA.002715505.1.protein:1889.58000,
(GB.GCA.002167825.1.protein:1743.61000,
(GB.GCA.002938255.1.protein:1579.05000,((GCA.902567835.1.AG-414-
E02.genomic:789.71000,(GCA.902515705.1.AG-325-E20.genomic:399.36200,
(GCA.902535465.1.AG-349-D08.genomic:230.39100,GCA.902609855.1.AG-917-

K20.genomic:230.39100)Node_598:168.97100)Node_597:390.34800)Node_596:8
6.48870,((GCA.902576855.1.AG-426-
E17.genomic:381.55300,RS.GCF.001180105.1.protein:381.55300)Node_609:16
1.30100,(GCA.902574695.1.AG-422-N15.genomic:464.15800,
((GCA.902617415.1.AG-892-F15.genomic:252.26900,
(GCA.902605865.1.AG-461-L09.genomic:104.50700,GCA.902583075.1.AG-912-
G22.genomic:104.50700)Node_603:147.76200)Node_602:140.32800,
(GCA.902569405.1.AG-414-L04.genomic:314.16700,(GCA.902579985.1.AG-430-
B09.genomic:175.20900,(RS.GCF.001179925.1.protein:112.06500,
(GCA.902555725.1.AG-390-D15.genomic:94.25140,
(RS.GCF.000701385.1.protein:51.74680,RS.GCF.000504225.1.protein:51.746
80)Node_608:42.50450)Node_607:17.81350)Node_606:63.14400)Node_605:138.
95800)Node_604:78.43030)Node_601:71.56090)Node_600:78.69660)Node_599:3
33.34400)Node_595:702.85200)Node_594:164.56300)Node_593:145.96500)Node
_592:278.87700,((GB.GCA.002695035.1.protein:1957.90000,
(GB.GCA.900551585.1.protein:1850.94000,GB.GCA.002389385.1.protein:1850
.94000)Node_644:106.95700)Node_643:102.78400,
((GB.GCA.002694385.1.protein:1743.29000,
(RS.GCF.003173035.1.protein:1626.50000,
(GB.GCA.003248505.1.protein:1520.88000,
(((GB.GCA.003164925.1.protein:1072.40000,GB.GCA.002307725.1.protein:10
72.40000)Node_621:174.30300,
(RS.GCF.000007125.1.protein:1089.10000,GB.GCA.002715765.1.protein:1089
.10000)Node_620:157.60200)Node_619:110.12300,
(RS.GCF.000170875.1.protein:351.35200,
(RS.GCF.000152965.1.protein:238.60500,
(RS.GCF.000014045.1.protein:124.79800,RS.GCF.005887615.1.protein:124.7
9800)Node_618:113.80700)Node_617:112.74700)Node_616:1005.48000)Node_61
5:164.05400)Node_614:105.61400)Node_613:116.79500)Node_612:167.17500,
((GB.GCA.001767835.1.protein:1731.65000,
(RS.GCF.000283655.1.protein:1586.39000,
(RS.GCF.002723915.1.protein:1293.35000,GB.GCA.004297265.1.protein:1293
.35000)Node_642:293.03800)Node_641:145.25900)Node_640:83.26410,
((GB.GCA.002724635.1.protein:1568.34000,
(GB.GCA.002709075.1.protein:1417.11000,GB.GCA.002501105.1.protein:1417
.11000)Node_625:151.23000)Node_624:164.54800,
((GB.GCA.002709955.1.protein:1524.91000,RS.GCF.900100655.1.protein:152
4.91000)Node_639:136.10700,(RS.GCF.900102465.1.protein:1405.09000,
((GB.GCA.003278095.1.protein:277.22300,(GCA.902619615.1.AG-892-
O07.genomic:151.37600,(GCA.902623995.1.AG-893-
P05.genomic:112.07100,GCA.902625215.1.AH-287-
L13.genomic:112.07100)Node_638:39.30530)Node_637:125.84600)Node_636:78
1.75700,((GCA.902625725.1.AH-287-O21.genomic:628.05900,
(GB.GCA.002387605.1.protein:505.14700,
(RS.GCF.000238815.1.protein:253.90300,GB.GCA.002348585.1.protein:253.9
0300)Node_632:251.24400)Node_631:122.91200)Node_630:214.15400,
(GB.GCA.002457745.1.protein:592.95600,
(GB.GCA.004212735.1.protein:484.16200,
(RS.GCF.000024465.1.protein:375.73900,GB.GCA.003282865.1.protein:375.7
3900)Node_635:108.42300)Node_634:108.79400)Node_633:249.25800)Node_629

:216.76700)Node_628:346.10500)Node_627:255.93000)Node_626:71.87400)Node_623:82.02440)Node_622:95.55410)Node_611:150.21200)Node_610:107.77600)Node_591:213.47900)Node_590:276.32600)Node_589:105.41700)Node_54728992663.26:215.06500)Node_546:114.56800)Node_545:319.89200)Node_544:293.71800)Node_543:149.93100)Node_542:78.39540)Node_541:36.90540)Node_540:149.41600)Node_539:86.95750)Node_5384331.65:10.14590)Node_537:437.84500);
