## Supplemental File 3-5_7-8 for "A Timeline of Bacterial and Archaeal Diversification in the Ocean"

| Parameter | Effective Size | Relative difference |
|---|---|---|
| **Loglik** | **7** | **0.16** |
| **Length** | **12** | **0.30** |
| **Sigma** | **21** | **0.16** |
| **Mu** | **37** | **0.09** |
| **Meanrates** | **19** | **0.16** |
| Scale | 19 | 0.57 |
| Alpha | 26 | 0.92 |
| **Nmode** | **70** | **0.29** |
| **Stat** | **8** | **0.37** |
| Statalpha | 20 | 0.94 |
| **Kappa** | **362** | **0.13** |
| Allocent | 10 | 0.51 |

Number of cycles = 92412; burn-in = 250; sampling = 2 cycles; 4 chains

**Supplemental File 4**

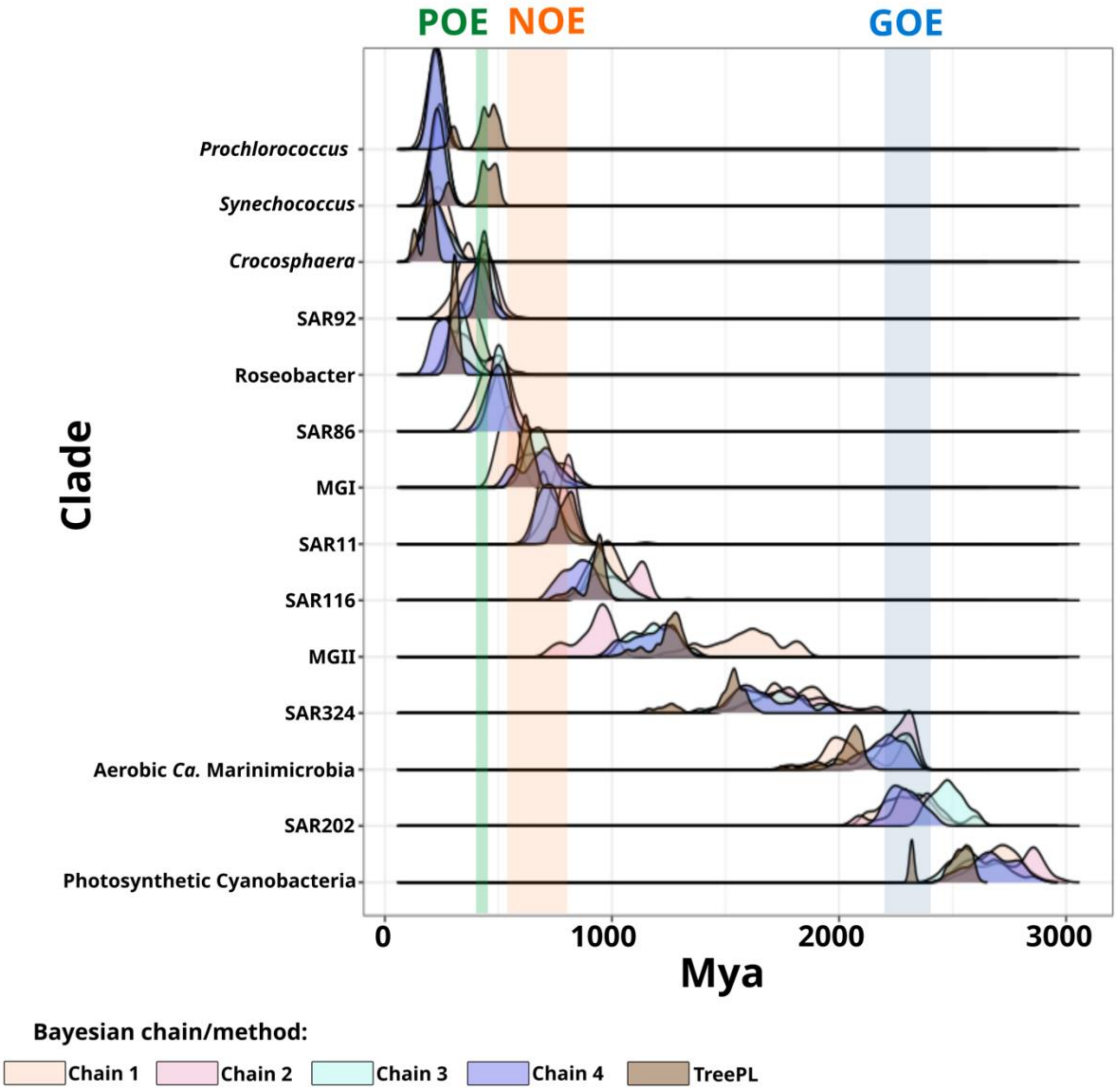

**Supplemental File 5**

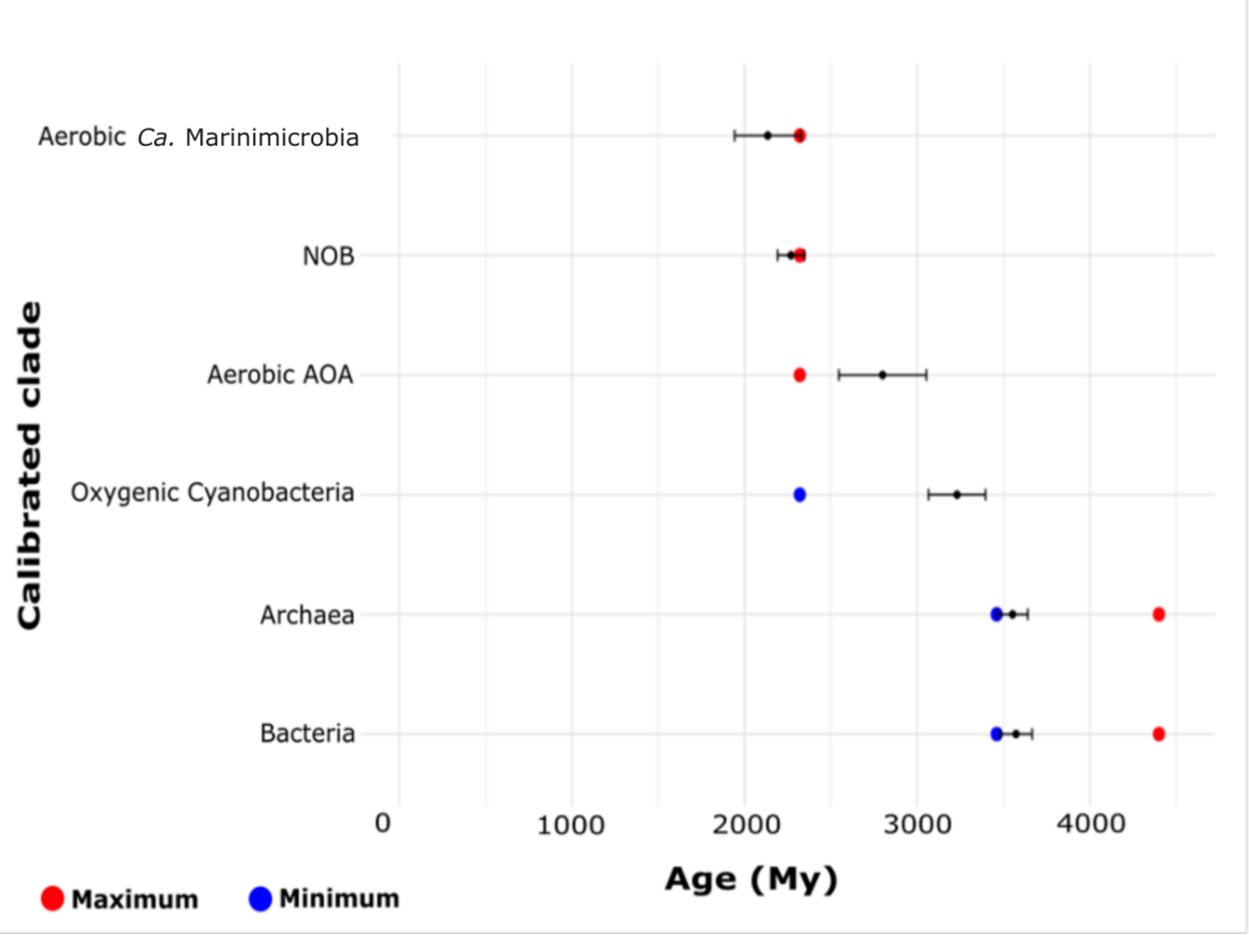

**Supplemental File 7**

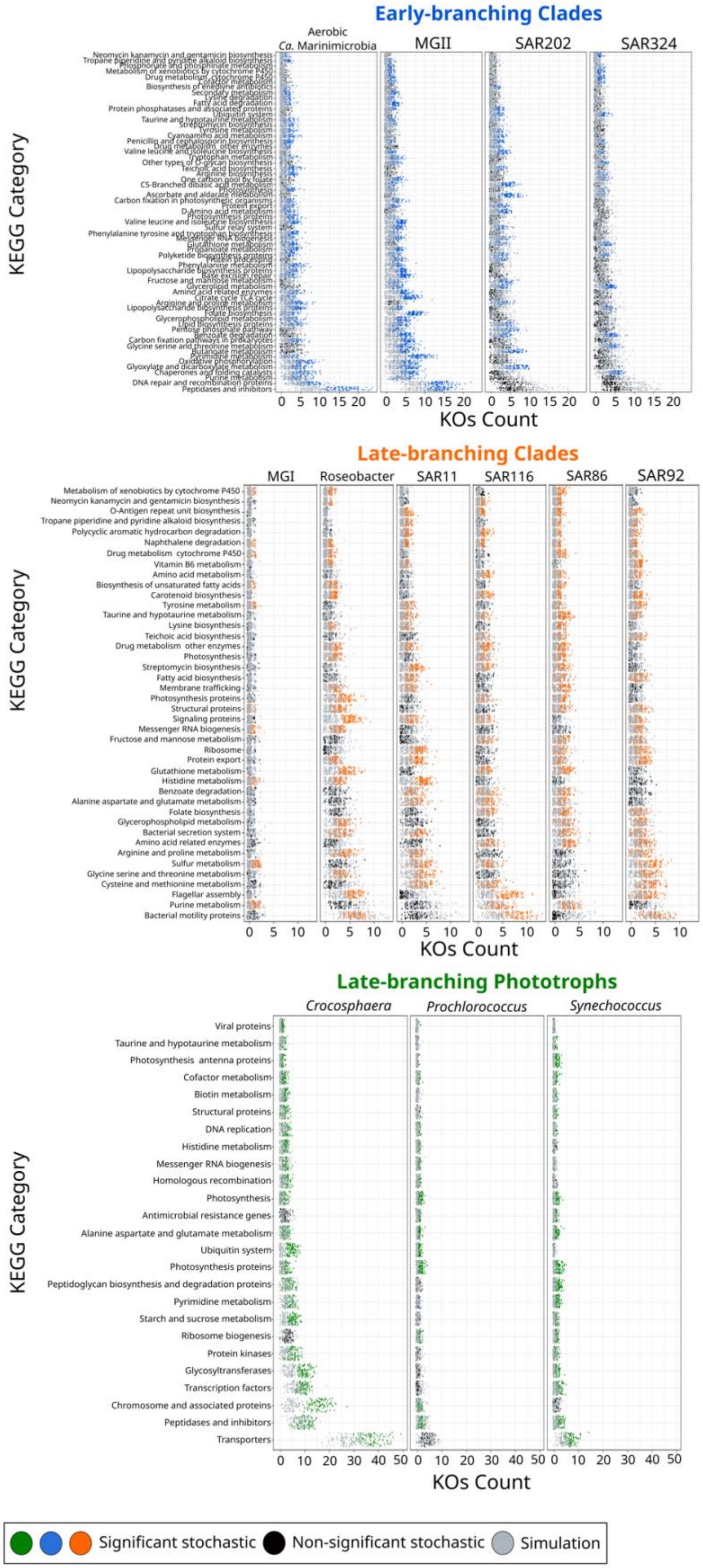


**Supplemental File 8**

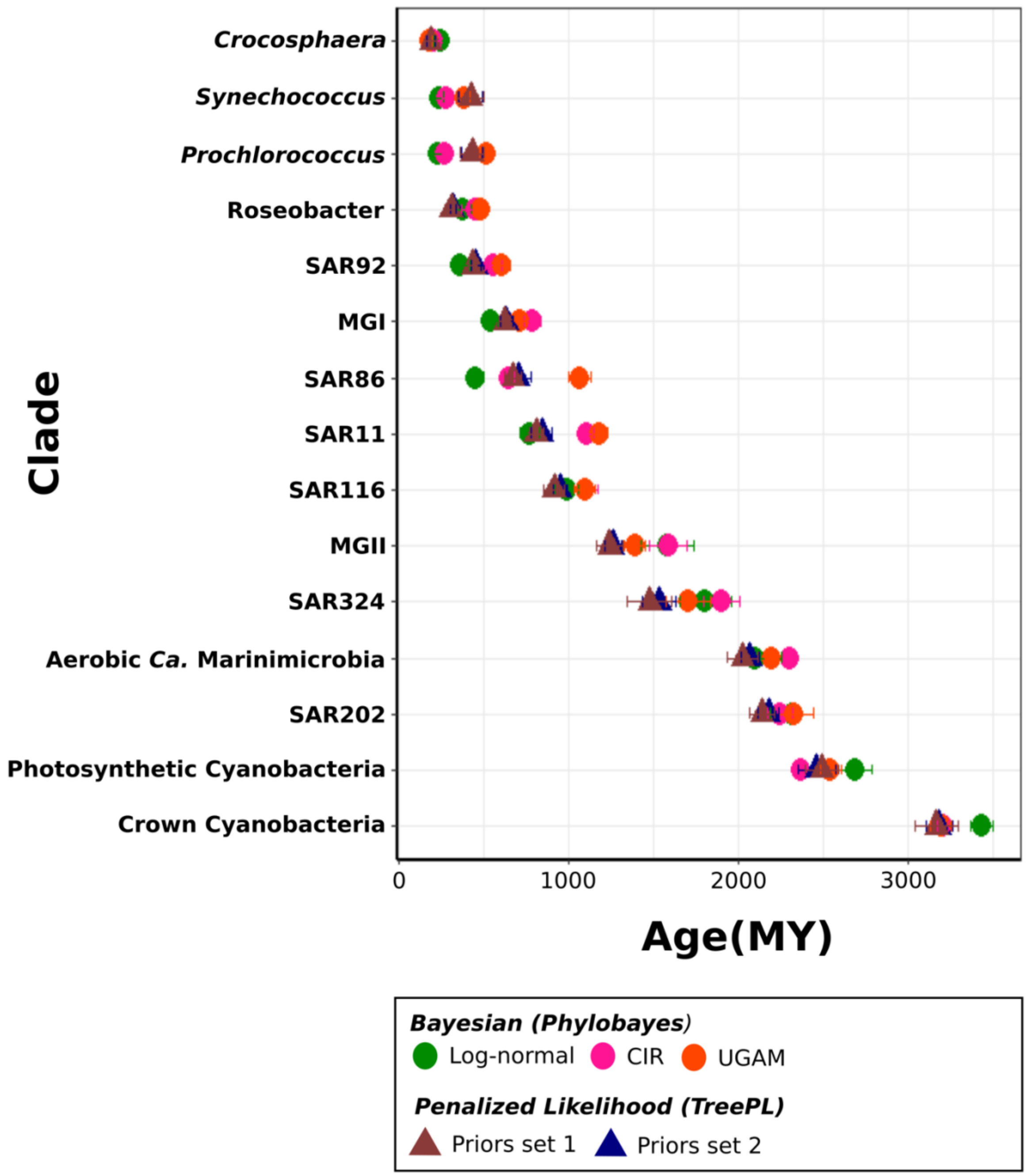
Crocosphaera
Synechococcus
Prochlorococcus
Roseobacter
SAR92
MGI
SAR86
SAR11
SAR116
MGII
SAR324
Aerobic Ca. Marinimicrobia
SAR202
Photosynthetic Cyanobacteria
Crown Cyanobacteria
Clade
0
1000
2000
3000
Age(MY)
Bayesian (Phylobayes)
Log-normal
CIR
UGAM
Penalized Likelihood (TreePL)
Priors set 1
Priors set 2
